# Charge segregation in the C-terminal linker of FtsZ enables Z-ring scaling with prokaryotic cell width

**DOI:** 10.1101/2025.11.25.690400

**Authors:** Marko Kojic, Daniela Megrian, Martin Loose

## Abstract

Bacterial cytokinesis is orchestrated by the Z-ring, a cytoskeletal structure formed by treadmilling filaments of the tubulin-like GTPase FtsZ. During assembly, filament curvature must match the cell diameter, yet how curvature is encoded in FtsZ remains unclear. Using *in vitro* reconstitution with purified proteins and supported lipid bilayers, we identify FtsZ’s intrinsically disordered C-terminal linker (CTL) as a curvature tuner that determines the Z-ring diameter. Increasing CTL charge segregation (κ) progressively reduces Z-ring diameter without altering treadmilling dynamics, monomer turnover, or GTPase activity, indicating that κ modulates filament curvature rather than polymerization kinetics. Linker-dilution experiments, in which wild-type FtsZ is mixed with a CTL-deletion mutant (ΔCTL), further show that inter-CTL electrostatic interactions straighten filaments, while weakening these contacts increases curvature and compacts rings.

To test whether this mechanism is conserved, we analyzed 4,603 prokaryotic FtsZ sequences and found that CTL charge segregation quantified by κ – but not CTL length, fraction of charged residues, or net charge per residue – correlates with cell width: at low κ, cells show a broad range of widths, whereas high κ values exclude wide cells. Co-expression of ΔCTL variants *in vivo* perturbs Z-ring positioning and impairs growth, consistent with a curvature mismatch and defects during assembly. Together, these results suggest a simple evolutionary mechanism in which CTL charge patterning encodes intrinsic filament curvature, enabling Z-ring geometry to scale with cell diameter and linking protein sequences to bacterial morphology.

## Introduction

Cell division is a fundamental biological process. In most prokaryotes, division proceeds by binary fission, in which the cell constricts at the middle to generate two identical daughter cells. This process is initiated and organized by the tubulin-like GTPase FtsZ, which forms a dynamic, ring-like cytoskeletal scaffold of treadmilling filaments^1–4^. This structure, termed the Z-ring, coordinates the stepwise recruitment of additional components to form the divisome, a multi-protein machinery that controls cell wall remodeling and cell constriction^5^.

FtsZ polymerization *in vivo* is under tight control^6^. Spatial regulators, including the Min system^6,7^, and nucleoid occlusion^8^, restrict FtsZ polymerization away from the poles and unsegregated chromosomes, respectively. FtsZ-associated proteins such as ZapA help shape Z-ring architecture by crosslinking filaments^9^, while treadmilling facilitates removal of misaligned filaments at mid-cell^10^. Recent simulations of treadmilling filaments suggest that filament curvature plays an important role in the perpendicular alignment of filaments to the long cell axis^10^. However, the determinants of FtsZ filament curvature and its relationship to cellular geometry remains mechanistically unresolved.

Curvature sensing by cytoskeletal elements is known to play an important role in shaping bacterial cells. The actin-like proteins MreB^11^ and FtsA^12^ form filaments that act as scaffolds for the assembly of protein complexes responsible for cell-wall remodeling during elongation and division, respectively. *In vitro,* their filaments exhibit small intrinsic radii of curvature of ∼200 nm^11^ in the case of MreB and ∼16 nm^12^, for FtsA. This has led to the suggestion that they align perpendicular to the long cell axis *in vivo* by binding to membrane regions of highest negative Gaussian curvature, thereby promoting circumferential peptidoglycan synthesis perpendicular to the long cell axis.

In contrast, purified FtsZ form filaments that exhibit a wide spectrum of curvatures depending on the experimental conditions: FtsZ mini-rings imaged by electron microscopy (∼25 nm radius)^13^; arcs and rings on mica surfaces imaged by AFM (150 to 300 nm)^14^; and larger rings in solution in the presence of ZapD (outer ring diameter ∼500 nm)^15^. On supported lipid bilayers (SLBs), either when recruited by FtsA or as chimeras with YFP and a membrane targeting sequence (mts), FtsZ filaments organize into large-scale structures of rotating rings with diameters of around 1.25 µm, corresponding to the curvature of individual treadmilling filaments and comparable to the diameter of a typical *E. coli* cell^2,16,17^. As FtsZ filaments are flexible, they straighten at higher surface densities or in the presence of the crosslinker ZapA, due to increased packing density and enhanced lateral contacts^9,18^. The close match between the curvature of membrane-bound FtsZ filaments *in vitro* and cell geometry suggests that filament curvature contributes to the correct orientation of protofilaments during initial Z-ring assembly, while the observed flexibility of FtsZ filaments is important to allow for the progressive constriction of the Z-ring during cell division. Given the diversity of bacterial shapes and sizes^19,20^, this raises the question of how FtsZ’s intrinsic filament curvature is established and tuned across species to match cellular geometry.

As steric repulsion between intrinsically disordered regions (IDRs) of proteins is known to contribute to the forces that deform membranes^21,22^, we set out to test the hypothesis that the biophysical properties of FtsZ’s intrinsically disordered C-terminal linker (CTL) could set filament curvature. Using *in vitro* reconstitution on supported lipid bilayers (SLBs) and quantitative fluorescence microscopy, we show that charge segregation in the CTL controls Z-ring geometry without altering polymerization dynamics or GTPase activity. Comparative analyses across 4603 prokaryotic FtsZ sequences further reveal an evolutionary coupling between CTL electrostatics and cellular geometry.

Together, our results indicate that the CTL’s charge distribution provides a sequence-encoded mechanism that tunes intrinsic filament curvature, allowing Z-ring geometry to scale with cell diameter and ensuring robust division across diverse bacterial morphologies.

## Results

### Evolutionary analysis reveals that cell width scales with charge segregation in CTL

On SLBs, *E. coli* FtsZ polymerizes into treadmilling filaments with an intrinsic curvature that matches that of the *E. coli* cell^2,5,18^. Because bacterial cell diameters vary by more than an order of magnitude^23^, we asked if FtsZ encodes structural information that allows its filament curvature to scale with cell width.

FtsZ’s globular domain is known to be highly conserved^24^, while the CTL is known to be intrinsically disordered^13,25^ as well as highly variable in length and sequence^24^ (**Fig. 1a**). Given the low degree of conservation and known properties of IDRs to tune protein-protein interactions^26^, and to contribute to membrane curvature sensing and generation^21,22^, we hypothesized that any scaling of filament curvature with cell diameter could be encoded within the CTL and depend on its biophysical properties.

**Fig. 1.**
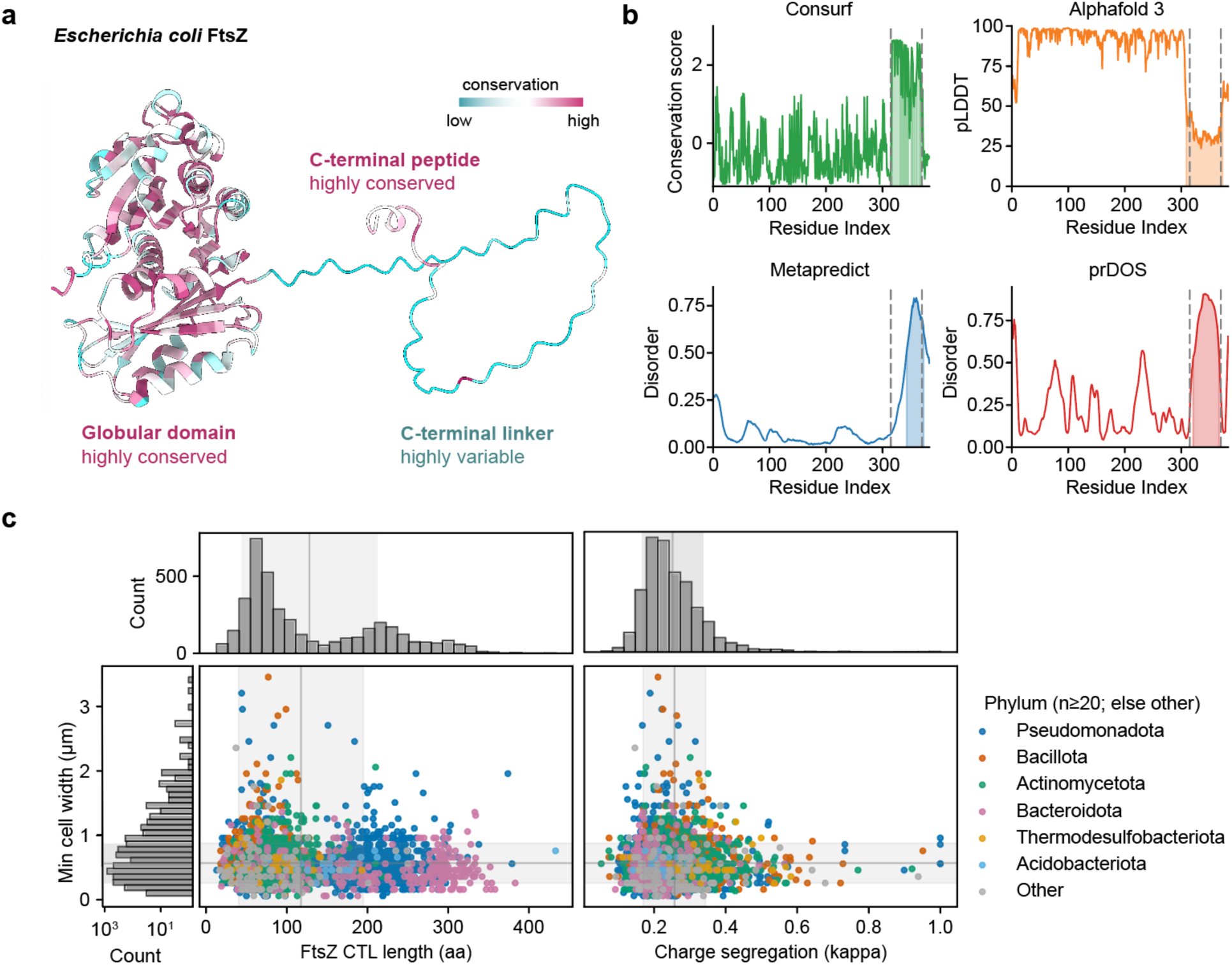
Relationship between the properties of the CTL and cell width. **(a)** Multiple sequence analysis (MSA) scores are mapped onto the predicted structure of *E. coli* FtsZ. **(b)** Different approaches on identifying an IDR region in *E. coli* FtsZ: sequence conservation (Consurf), structure prediction (AlphaFold3), prediction of structural disorder (Metapredict and PrDOS). Gray dotted lines represent residue positions defining the region of CTL as denoted in the literature. **(c)** Relationship between cell widths and the length of CTL (left) or CTL’s charge segregation (right) analyzed using Metapredict-identified CTLs and Sparrow to calculate the CTL length and charge segregation (k) values. Different colors represent different phyla. Histograms represent the distribution of CTL lengths (upper left), charge segregation (upper right) and cell widths (left). Gray line and gray-shaded area represent the mean and standard deviation of the corresponding parameters.

To identify any potential correlation between the sequence of FtsZ and cell dimensions of a given organism, we compiled 4603 FtsZ sequences from bacterial and archaeal species with cell widths annotated in the BacDive database^27^. Reported minimal cell widths ranged from 0.1 µm to > 3.5 µm (mean ± SD = 0.606 ± 0.308 µm). Alignment of the FtsZ sequences of this dataset (**Supplementary Information**) and a Consurf^28^ analysis based on *E. coli* FtsZ (**Fig. 1b**), demonstrate the high conservation of the GTPase domain and great variability between residues 315 and 365, the previously reported boundaries of the CTL in this protein^29^. We used three independent computational approaches to identify the CTL in the sequences of our dataset: Metapredict, a deep learning tool that predicts disorder directly from sequence by reproducing consensus disorder profiles^30^; PrDOS, which integrates sequence-derived machine learning features with evolutionary evidence from homologous structures^31^ and AlphaFold2, which infers disorder from low-confidence regions in its structure-based pLDDT scores^32^. All three predictors confirm the disordered character of *E. coli* FtsZ’s CTL, despite some differences in identifying the boundaries of CTL (**Fig. 1b**).

Because the length of IDRs was found to modulate steric and entropic interactions between proteins^29,33–35^, we were first interested in identifying any correlation between CTL length and cell size. Our analysis showed that CTL lengths ranged from 13 to 433 amino acids (mean ± SD = 128.5 ± 84.6 amino acids), exhibiting a bimodal overall distribution. This is mainly driven by Bacteroidota and Pseudomonadota (**Fig. 1c, Fig. S1a**), where Bacteroidota sequences showed a prominent peak around ∼295 aa, whereas Pseudomonadota displayed two distinct peaks, one near ∼75 aa and another around ∼200 aa, reflecting considerable variability within this phylum. However, CTL length showed no substantive correlation with mean cell width (Spearman ρ = −0.081). Cells with diameters around the mean (∼0.6 µm) contained FtsZ CTLs spanning a wide range (16–382 aa), similar to wider cells (> 1.5 µm), which also exhibited broad CTL lengths (37– 374 aa). Together, these findings indicate that cell diameter and consequently the initial Z-ring diameter does not scale with the length of the FtsZ CTL.

Next, we used SPARROW, a deep-learning model for predicting properties of IDRs, to calculate for each CTL its radius of gyration, fraction of charged residues (FCR), net charge per residue (NCPR), and charge segregation (κ)^36^. These parameters describe the electrostatic organization and compaction tendencies of an IDR, as well as their tendencies for intermolecular interactions^37^. Neither FCR nor NCPR correlated meaningfully with cell width (Spearman ρ ≈ −0.107 and 0.191, respectively; **Fig. S1b**). By contrast, we found that charge segregation (κ) was associated with cell width. Small cells (width = 0.1–1.0 µm) spanned nearly the full positive κ range (0.0407-1.000, mean ± SD = 0.25 ± 0.09), whereas large cells (> 1.5 µm) were restricted to a narrow low-κ interval (0.11–0.40, mean ± SD = 0.24 ± 0.06). Although the mean κ values for large and small cells were nearly identical, and the overall correlation between κ and cell width was weak and not significant (Spearman ρ = 0.02), wide cells were absent at high κ, while low κ allows for a wide distribution of cell width. We found similar results for IDRs identified using all three approaches (**Fig. S1a,b**). Together, these observations reveal a distinct “forbidden region” in the width–κ parameter space (**Fig. 1c**) and suggest that electrostatic patterning plays a role in constraining the geometry of the Z-ring.

Interestingly, AlphaFold2 structure predictions for the FtsZ sequences in our dataset further revealed that, in a subset of sequences, the CTL of FtsZ can adopt a partially ordered conformation. In these sequences, the CTL has continuous stretches of high-confidence residues (pLDDT > 70) with predicted α-helical elements and increased sequence conservation, suggesting structural constraints likely associated with species-specific molecular interactions. This feature was most prevalent in Bacteroidota (533 of 1,013 sequences), followed by Pseudomonadota (106 of 1,781) and Methanobacteriota (49 of 54) (**Fig. S1c**), These observations indicate that extensive intrinsic disorder is a dominant feature of the CTL across homologs, with specific departures from complete disorder arising in specific lineages, potentially reflecting additional species-specific interactions. These cases represent a minority within the dataset and do not alter the overall relationship between CTL electrostatic patterning and cell width described above.

### The length of the CTL does not affect Z-ring diameter *in vitro*

Our comparative analysis suggested that properties of the CTL might scale with the cell diameter. However*, in vivo*, the size of the Z-ring is limited by the geometry of the cell and accessory proteins, such as ZapA^9^ and ZapD^15^, which can modify the curvature of FtsZ filaments. In addition, electron microscopy studies found evidence that the CTL influences lateral interaction between protofilaments^33–35,38^, but these studies could not reveal the dynamic emergent properties of dynamic, treadmilling filaments on membrane surfaces. To understand how the properties of the CTL affect the dynamic large-scale organization of FtsZ filaments, we took advantage of our previously established *in vitro* assay. Here, purified FtsZ and FtsA proteins are incubated on SLBs, where FtsZ polymerizes into treadmilling filaments that further organize into rotating rings (**Fig. 2a**)^2^. In this experiment, we have full control of the biochemical composition of the assay as well as the FtsZ linker sequence. In addition, the absence of spatial boundaries allows to reveal the intrinsic curvature of FtsZ filaments.

**Fig. 2.**
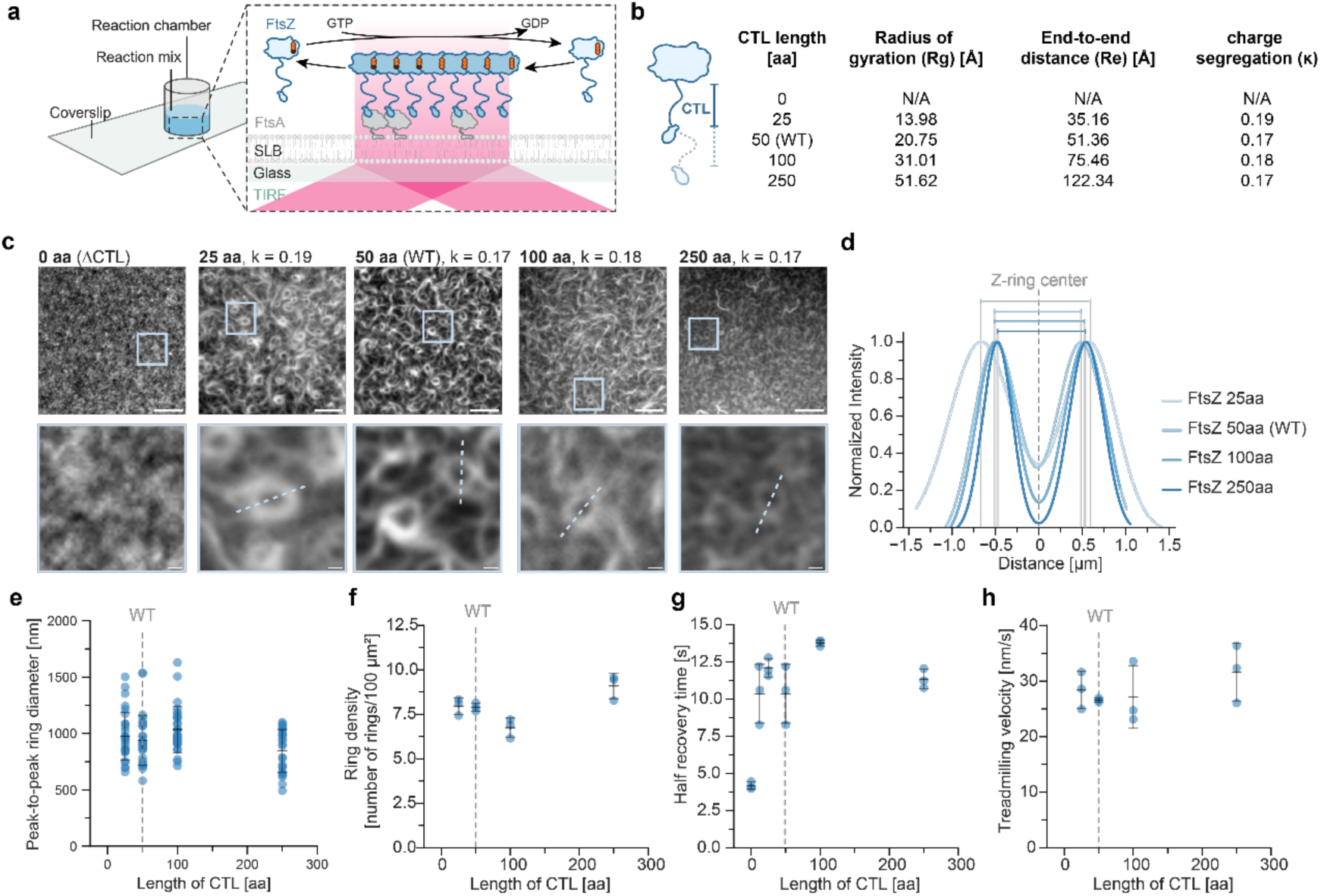
CTL length does not change FtsZ filament networks. **(a)** Schematic representation of the experimental set-up. **(b)** Biophysical parameters of different FtsZ CTL length mutants: CTL length (aa), Radius of gyration (Rg) (Å), End-to-end distance (Re) (Å), and charge segregation (κ). **(c)** TIRF micrographs of fluorescently labeled FtsZ CTL length mutants (1.5 µM) on SLBs recruited by FtsA (0.2 µM). Scale bar: 5 µm. Blue squares represent regions of the corresponding micrograph shown at 10x magnification shown below. Scale bar: 500 nm. **(d)** Normalized fluorescent intensity profiles along the dotted lines drawn in c. **(e)** Analysis of peak-to-peak ring diameter (nm) in respect to the length of CTL. **(f)** Analysis of ring density (number of rings/100 µm^2^), **(g)** half recovery time (s) and **(h)** treadmilling velocity (nm/s) in respect to the length of CTL. **(e-h)** Dotted gray line represents the value of FtsZ WT.

We first asked whether altering CTL length alone, while keeping charge segregation approximately constant, could affect FtsZ filament curvature and consequently the Z-ring diameter. To test this hypothesis, we generated and purified *E. coli* FtsZ variants with CTL lengths of 0 aa (ΔCTL), which still included the C-terminal peptide essential for FtsA binding^39^, 25 aa, 50 aa (WT), 100 aa, and 250 aa, maintaining κ ≈ 0.17–0.19 (**Fig. 2b; Supplementary Movie 1**). FtsZ versions with CTLs of 100 aa and 250 aa were designed using the *A. tumefaciens* CTL sequence as described previously^38^. Despite some variation among three different methods, disorder-prediction analyses confirmed that all constructs except FtsZ ΔCTL retained a C-terminal IDR (**Fig. S2a-e)**. Each fluorescently labeled FtsZ variant at the concentration of 1.5 µM was incubated with 0.2 µM FtsA on SLBs, which supports robust formation of rotating rings for wild-type FtsZ^18^.

FtsZ ΔCTL failed to form filament patterns (**Fig. 2c; Supplementary Movie 1**), indicating that the CTL is required for large-scale self-organization. Increasing the concentration of this mutant up to 6 µM did not restore pattern formation (**Fig. S3a; Supplementary Movie 2**). Filament turnover of FtsZ ΔCTL was increased as indicated in reduced half recovery time (**Fig. S3b; Supplementary Movie 3**), while showing a modest decrease in GTPase activity (**Fig. S3c**), suggesting that filaments FtsZ ΔCTL cannot be recruited to membranes robustly, possibly to due steric effects and the missing flexibility of the IDR.

In contrast, all tested FtsZ variants containing a CTL formed membrane-bound networks of treadmilling filaments (**Fig. 2c**), which organize into polar streams and rotating rings as described for *E. coli* FtsZ WT previously^2,18^. Quantitative image analysis revealed that outer and inner diameters of FtsZ rings, their peak-to-peak distances, and ring densities were independent of CTL length (**Fig. 2d-f, Fig. S2f-h**). Although a small but statistically significant difference was detected between the 25 aa and 250 aa variants (**Fig. 2d**: (974.63 ± 211.53 nm and 845.46 ± 189.09 nm, respectively; p = 0.03), no systematic trend with increasing length was observed. Likewise, fluorescence recovery after photobleaching (FRAP) experiments showed similar recovery kinetics across all variants (**Fig. 2g; Supplementary Movie 3**), and treadmilling velocities and GTP hydrolysis rates were indistinguishable from wild type (**Fig. 2h, Fig. S2i)**, suggesting similar filament lengths^40^.

These results agree with previous *in vivo* observations that the CTL is indispensable for Z-ring formation^29,41^ and that both *E. coli* and *B. subtilis* tolerate a wide range of CTL lengths without division defects^29,41^. They also show the CTL length alone does not set filament curvature and thereby the Z-ring diameter. The intrinsic curvature of the FtsZ filament must therefore be determined by other features.

### Charge segregation pattern within the CTL determines the Z-ring diameter *in vitro*

Given that the comparative analysis between cell width and charge segregation (κ) indicated a strong relationship, we aimed to address directly how κ affects the Z-ring size *in vitro*. As noted by Cohan *et al.*^41^, κ is most informative when comparing sequences with identical composition and length. We therefore designed a set of FtsZ variants in which only the patterning of charged residues in the 50-residue CTL, but not sequence length or composition, was altered. We constructed a series of FtsZ variants with linkers spanning a broad range of charge segregation values (κ = 0.13, 0.17 (WT), 0.34, 0.53, 0.85, 1.00), all predicted to be intrinsically disordered (**Fig. 3a, Fig. S4a–f**). These FtsZ variants were fluorescently labeled and their polymerization and self-organization on membranes reconstituted as before.

**Fig. 3.**
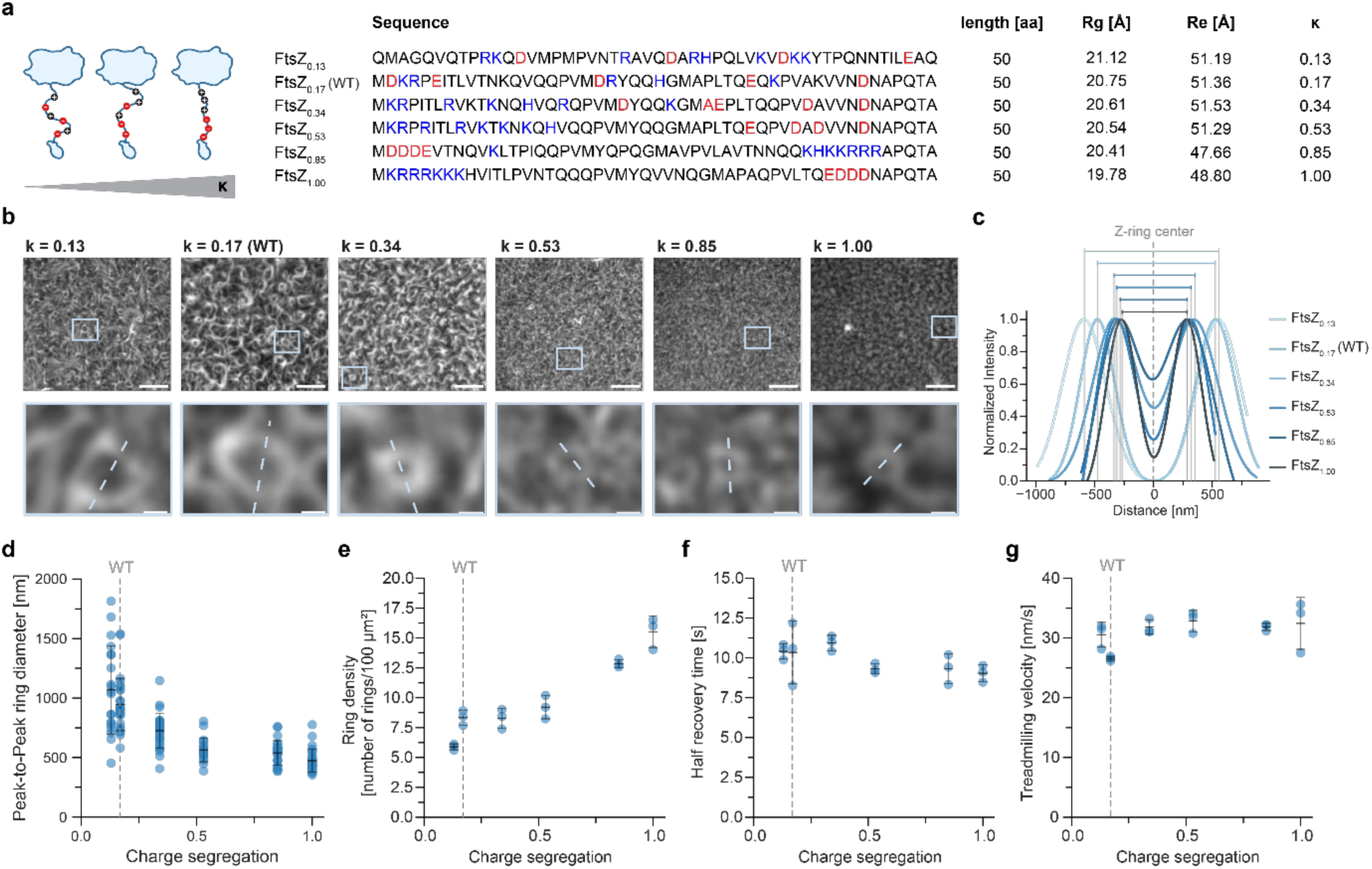
Charge segregation in FtsZ’s CTL determines the Z-ring diameter *in vitro*. **(a)** Schematic representation of charge segregation patterns, CTL sequences of FtsZ charge segregation mutants used in this study, and their biophysical parameters. **(b)** TIRF micrographs (top) of fluorescently labeled FtsZ CTL charge segregation mutants (1.5 µM) on SLBs recruited by FtsA (0.2 µM). Scale bar: 5 µm. Blue squares represent regions of the corresponding micrograph shown at 10x magnification shown below. Scale bar: 500 nm. **(c)** Normalized fluorescent intensity profiles along the dotted lines drawn in b. **(d)** Analysis of peak-to-peak ring diameter (nm), **(e)** ring density (number of rings/100 µm2), **(f)** half recovery time (s) and **(g)** treadmilling velocity (nm/s) in respect to the charge segregation in CTL. **(d-g)** Dotted gray line represents the value for FtsZ WT.

At low κ values, FtsZ formed extended ring-like patterns with diameters comparable to wild-type assemblies (**Fig. 3b; Supplementary Movie 4**). As κ increased, the peak-to-peak ring diameter decreased progressively from 1.07 ± 0.37 to 0.47 ± 0.10 µm (**Fig. 3c,d**), accompanied by an increase in ring density (**Fig. 3e**). Notably, the two variants with κ = 0.85 and κ = 1.00 exhibited indistinguishable organization despite having inverted distributions of acidic and basic residues, underscoring that charge segregation strength rather than polarity dictates assembly behavior.

Despite these geometric changes, FRAP recovery kinetics, treadmilling velocities (**Fig. 3f,g; Supplementary Movie 5**), and GTP hydrolysis rates (**Fig. S4i**) remained unchanged, ranging from 9.04–10.95 s (FRAP recovery rates), 26.58–32.84 nm/s (treadmilling velocities) and 1.15–1.52 μM/min (GTP hydrolysis rates), indicating that the altered organization arises from differences in filament curvature rather than polymerization dynamics. Variants with low κ also displayed greater heterogeneity in ring diameters, echoing the broader distribution of cell widths observed among bacterial species with low CTL’s charge segregation (**Fig. 3d**). Co-assembly of the extreme variants FtsZ_0.13_ and FtsZ_1.00_ yielded mixed (Pearson’s correlation coefficient (PCC) = 0.94 ± 0.02), treadmilling networks with intermediate properties: slightly smaller ring diameters and increased density compared with FtsZ_0.13_ alone (**Fig. S5:** peak-to-peak ring diameter: 1.43 ± 0.47, 0.67 ± 0.13, 1.24 ± 0.28 (nm), and ring density: 5.90 ± 0.25, 15.53 ± 1.32, 7.93 ± 0.58 number of rings/100 µm^-2^; **Supplementary Movies 6,7**). Together, these results demonstrate that the electrostatic patterning of the CTL, quantified by κ, directly controls Z-ring geometry *in vitro*.

### Inter-CTL electrostatic interactions modulate Z-ring diameter *in vitro*

In the previous section, we found that increasing charge segregation (κ) within the FtsZ C-terminal linker (CTL) progressively reduced Z-ring diameter *in vitro*, suggesting that electrostatic patterning of the CTL directly influences filament curvature. We next asked how this effect arises at the molecular level.

Two mechanisms could, in principle, explain the κ-dependent scaling of the Z-ring size. First, intramolecular interactions within a single CTL might compact the disordered chain or modulate its contact with the globular domain, thereby altering filament curvature. Alternatively, intermolecular interactions between CTLs on adjacent subunits within an FtsZ filament could mediate attractive or repulsive forces that straighten or bend the polymer.

To distinguish between intra- and intermolecular interactions, we designed a “dilution” experiment in which CTL–CTL contacts are selectively weakened by increasing the averaged distance between consecutive CTLs within the FtsZ filament. This was achieved by mixing wild-type, fluorescent FtsZ (with intact CTL) with FtsZ ΔCTL **(Fig. 4a**). If inter-CTL interactions tune filament curvature, reducing the density of CTLs in a filament should lead to either more curved or straighter filaments *in vitro*. In contrast, if curvature is dictated primarily by intramolecular interactions, then reducing CTL density should not alter filament curvature, allowing us to discriminate between intra- and intermolecular mechanisms.

**Fig. 4.**
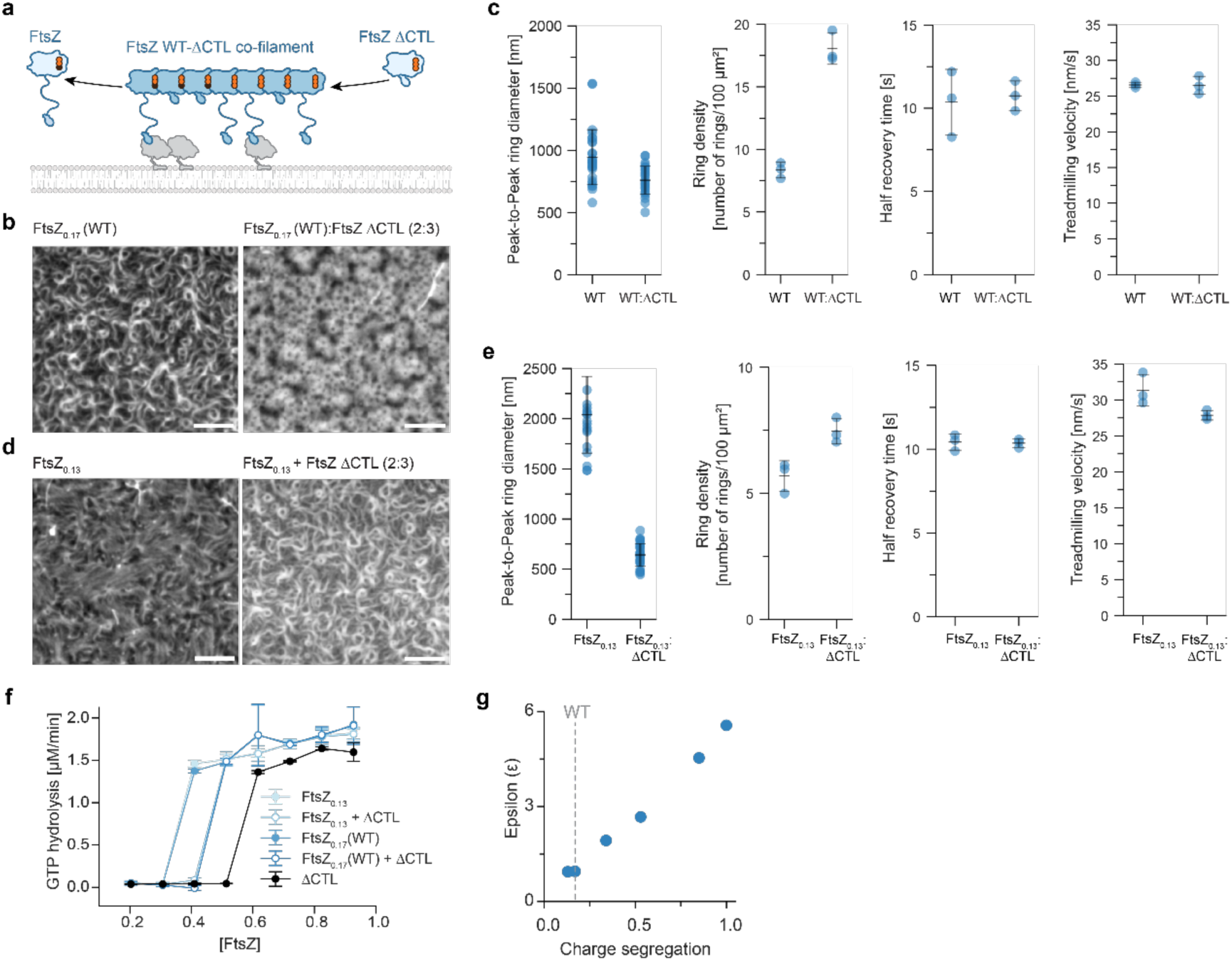
Inter-CTL electrostatic interactions modulate Z-ring diameter *in vitro*. **(a)** Schematic representation of the experimental set-up. **(b)** TIRF micrographs of fluorescently labeled FtsZ_0.17_(WT) alone and in the presence of FtsZ ΔCTL (2:3; 1.5 µM) on SLBs recruited by FtsA. Scale bar: 5 µM. **(c)** Analysis of peak-to-peak ring diameter (nm), ring density (number of rings/100 µm^2^), half recovery time (s) and treadmilling velocity (nm/s) of b. **(d)** TIRF micrographs of fluorescently labeled FtsZ_0.13_ alone and in the presence of FtsZ ΔCTL (2:3; 1.5 µM) on SLBs recruited by FtsA. Scale bar: 5 µm. **(e)** Analysis of peak-to-peak ring diameter (nm), ring density (number of rings/100 µm^2^), half recovery time (s) and treadmilling velocity (nm/s) of c. **(f)** Critical GTPase concentration (Cc) calculation of FtsZ_0.17_(WT) and FtsZ_0.13_ alone and in the presence of FtsZ ΔCTL (2:3); and FtsZ ΔCTL alone. Cc is defined as a concentration of FtsZ at which the GTP hydrolysis rate significantly increased from the baseline. **(g)** Correlation between CTL’s epsilon (ε) and charge segregation values of the FtsZ CTL charge segregation mutants.

We found that co-assembly of FtsZ_0.17_ (WT) with FtsZ ΔCTL resulted in smaller Z-ring diameters, and greatly increased higher ring densities compared with pure wild-type protein (**Fig. 4b,c; Supplementary Movies 8,9**). A similar effect was observed when the low-κ variant FtsZ_0.13_ was mixed with FtsZ ΔCTL: ring diameter decreased, while ring density increased (**Fig. 4d,e; Supplementary Movies 10,11**). Additionally, when FtsZ_1.00_ was mixed with FtsZ ΔCTL, no Z-ring formation was observed, similarly to what FtsZ ΔCTL alone formed on SLBs (**Fig. S6a,b; Supplementary Movies 12,13)**.

To verify that the observed effects arise from co-assembly of FtsZ variants, we used Atto633-FtsZ_0.13_ and Alexa488-FtsZ ΔCTL in dual-color experiments. The two proteins co-localized (**Fig. S6c:** PCC = 0.77 ± 0.03; **Supplementary Movie 14**), indicating the formation of mixed filaments. The resulting Z-rings were smaller than those formed by FtsZ_0.13_ alone (1233 ± 0.268 and 874 ± 158 nm), and ring density increased, while filament turnover rate and treadmilling velocity remained unchanged, consistent with the predictions of the dilution experiment (**Fig. S6d-g; Supplementary Movie 15**).

Next, we hypothesized that if inter-CTL interactions are repulsive in wild-type FtsZ, reducing them should facilitate filament assembly and thereby lower the critical concentration for GTP hydrolysis. Conversely, if these interactions are attractive, reducing them should increase the critical concentration by weakening filament association. To test this, we performed concentration-dependent GTPase assays. We found that the critical concentration increased when FtsZ_0.13_ and FtsZ_0.17_ (WT) were mixed with FtsZ ΔCTL, from 0.4 to 0.5 µM, while FtsZ ΔCTL alone exhibited an even higher critical concentration of 0.6 µM. In addition, the critical concentration for GTP hydrolysis increased progressively with increasing charge segregation (κ) (**Fig. 4f, Fig. S7a,b**). These observations are consistent with CTL-mediated attractive interactions that are weakened in FtsZ_0.17_(WT)–ΔCTL co-filaments and in FtsZ CTL variants with higher κ values.

While Shinn *et al.*^42^ reported that deletion of the CTL enhanced GTP hydrolysis in *B. subtilis* FtsZ, we did not observe any increase in GTPase activity under our conditions, even at high protein concentrations (**Fig. S3c**). Moreover, treadmilling speed and subunit turnover rate remained unaffected across κ variants (**Fig. 3f,g**), supporting the conclusion that electrostatic patterning within the CTL modulates filament curvature without altering the intrinsic catalytic activity of FtsZ.

To further test our proposed mechanism, we calculated an intermolecular interaction parameter (ε) using the FINCHES framework^43^, which computes force-field interaction maps (“intermaps”) between intrinsically disordered peptides. Modeling pairs of CTLs aligned as in a filament revealed a strong linear relationship between ε and κ (**Fig. 4g, Fig. S8**): low-κ CTLs were weakly repulsive, whereas high-κ CTLs were increasingly repulsive. While this model captures only electrostatic repulsion and omits potential solvent-mediated effects, it reproduces the overall trend observed experimentally that increasing charge segregation weakens favorable CTL–CTL interactions. Thus, the computational analysis supports the view that inter-CTL electrostatics provide a mechanism for tuning filament curvature and Z-ring geometry.

### Dilution of inter-CTL interactions modulates the positioning of the Z-ring *in vivo*

To probe the functional importance of CTL-mediated attraction for Z-ring assembly inside cells, we expressed different FtsZ CTL charge segregation variants from replicative plasmids in *E. coli* MG1655, which still expresses native FtsZ. This setup allows ectopically expressed variants to co-polymerize with the endogenous protein, thereby effectively diluting native inter-CTL contacts, similarly to what we established in our *in vitro* set-up. When expressed ectopically from an arabinose-inducible promoter, both the highly charge-segregated mutant FtsZ_1.00_ and FtsZ ΔCTL caused a pronounced reduction in growth rate (**Fig. 5a and Fig.S9a and b**). In contrast, co-polymerization of native FtsZ with low κ variants or additional FtsZ_0.17_(WT) had only mild effects (**Fig. S9a and b**). The similar phenotypes of FtsZ_1.00_ and FtsZ ΔCTL are consistent with both proteins having weakened inter-CTL attraction, thereby perturbing filament organization and cell division.

**Fig. 5.**
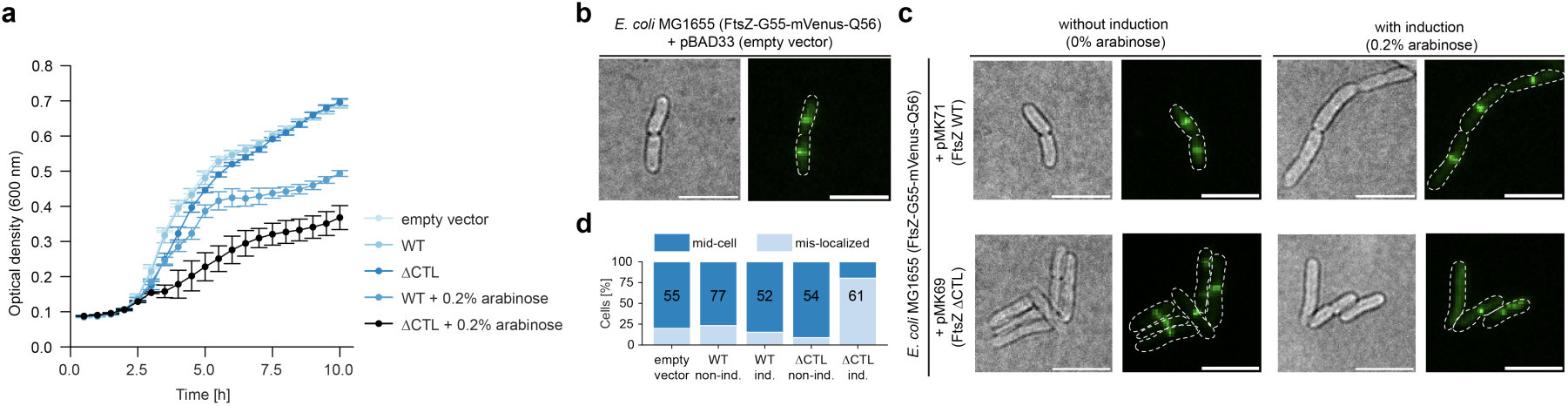
FtsZ WT/ΔCTL co-filaments alter Z-ring localization *in vivo*. **(a)** Growth curve analysis of *E. coli* MG1655 transformed individually with pBAD33 (empty vector), pMK71, pMK69, without or with induced expression (0.2% arabinose). **(b)** Direct Illumination Acquisition (DIA) and TIRF micrograph of *E. coli* MG1655 (FtsZ-G55-mVenus-Q56) transformed with pBAD33 (empty vector). Scale bar: 5 µm. **(c)** Top: DIA and TIRF micrographs of *E. coli* MG1655 (FtsZ-G55-mVenus-Q56) transformed with pMK71 (FtsZ WT) without (left) or with (right) induced expression (0.2% arabinose). Bottom: DIA and TIRF micrographs of *E. coli* MG1655 (FtsZ-G55-mVenus-Q56) transformed with pMK69 (FtsZ ΔCTL) without (left) or with (right) induced expression (0.2% arabinose). Scale bar: 5 µm. **(d)** Quantification of cells (%) with mis-localized Z-rings in a and b.

To examine how these perturbations affect Z-ring architecture, we used an *E. coli* strain carrying a chromosomally encoded, fully functional sandwich fusion of FtsZ (AV72; mVenus inserted between residues G55 and Q65)^44^ with wildtype CTL. While WT FtsZ formed tight rings at midcell, co-expression of FtsZ ΔCTL generated multiple Z-rings that failed to condense into a single structure. The fraction of cells with mis-localized or irregular Z-rings increased by nearly 60% relative to uninduced controls or cells expressing WT FtsZ ectopically (**Fig. 5b–d and Fig. S9c-e; Supplementary Movie 16**). These aberrant structures often appeared off-center or tilted relative to the cell axis, supporting the idea that reduced inter-CTL attraction impairs orthogonal Z-ring alignment within the confined cellular geometry.

Together, these *in vivo* experiments demonstrate interactions between CTLs are important for correct Z-ring positioning: when CTL–CTL interactions are weakened, division rings become misaligned or misplaced, leading to reduced growth rates. These findings show that the electrostatic organization of the CTL not only shapes filament curvature *in vitro* but also ensures to the spatial fidelity of cytokinesis *in vivo*.

## Discussion

FtsZ mediates cell division in most prokaryotes, including both Bacteria and Archaea. Present in the last universal common ancestor approximately 3.5 billion years ago, FtsZ has retained a highly conserved globular domain throughout evolution. For example, when comparing FtsZ from *B. subtilis* and *E. coli*, species whose last common ancestor is inferred to be the last bacterial common ancestor, the globular domains share 54% sequence identity and their crystallographic structures align with an RMSD of only 1.1 Å across 301 residue pairs^45,46^. This high conservation is particularly pronounced at the interface between monomers, where residues directly mediate the longitudinal contacts essential for polymerization^47^. Such strong evolutionary constraint on the globular domain is critical, as even minor perturbations to this region could disrupt longitudinal interactions, interfere with GTPase activity, or affect the conformational change of FtsZ essential for treadmilling dynamics^48^.

In contrast, the C-terminal linker (CTL) is subject to far fewer evolutionary constraints. As an intrinsically disordered region, it does not rely on specific residues to maintain a defined three-dimensional fold and can therefore vary widely in sequence, length, and physicochemical properties. This flexibility allows the CTL to evolve new functional features without compromising the essential polymerization behavior of the conserved globular domain.

Here, we uncover a biophysical mechanism by which the electrostatic properties of FtsZ’s CTL control filament curvature and, consequently, Z-ring geometry. The intrinsically disordered CTL offers a flexible and evolvable alternative for modulating curvature without compromising core filament properties. We find that weak attractive interactions between neighboring CTLs – in individual protofilaments and likely also within membrane-bound filament bundles – straighten filaments. In contrast, reducing these attractions increases curvature. This effect can be achieved either by increasing the charge segregation parameter (κ), which separates positive and negative residues into blocks, or by diluting full-length FtsZ with the ΔCTL mutant, thereby effectively increasing the distance between linkers (**Fig. 6**).

**Fig. 6.**
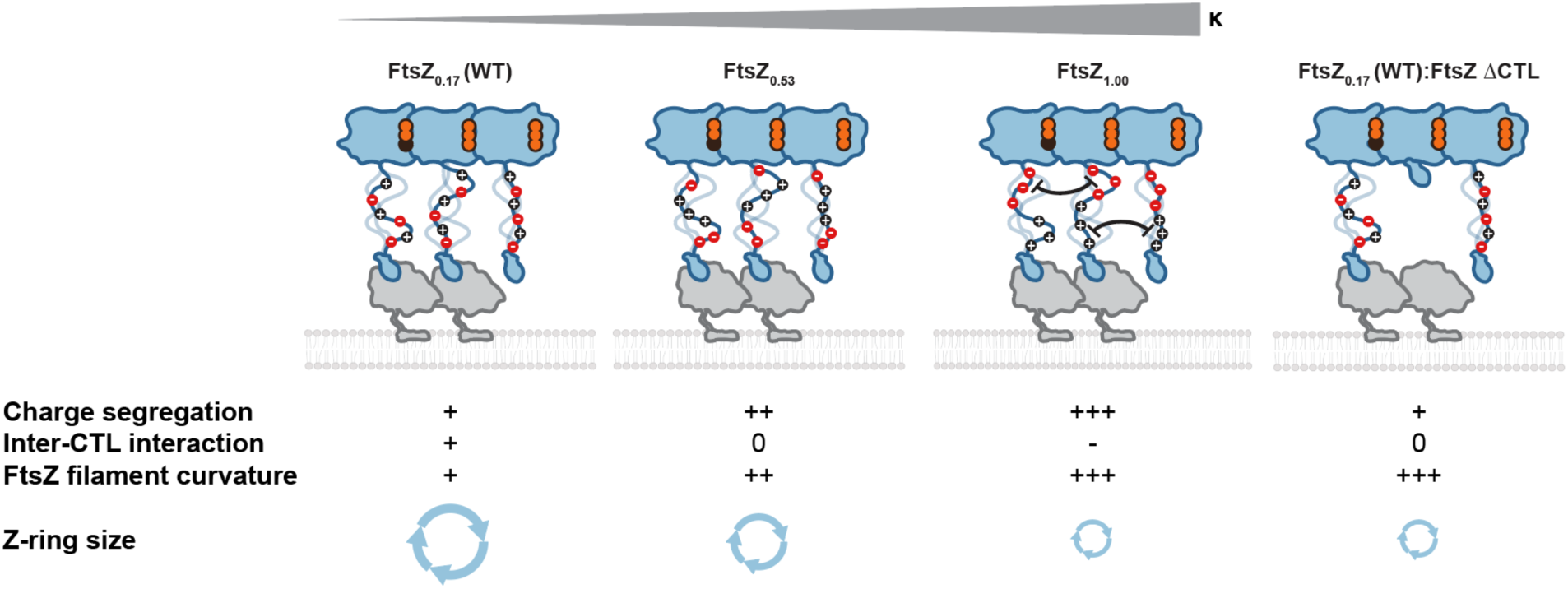
Proposed model for how CTL charge segregation tunes Z-ring architecture through modulation of inter-CTL electrostatics and filament curvature. Schematic illustrating the proposed mechanism by which increasing charge segregation within the FtsZ C-terminal linker (CTL) enhances inter-CTL electrostatic repulsion, leading to altered filament curvature and consequently the Z-ring size. Low κ FtsZs exhibit minimal repulsion and form low-curved filaments that assemble into large Z-rings. Intermediate and high κ FtsZs enforce stronger inter-CTL repulsion, increased filament curvature, and small Z-ring diameters. Our dilution experiments (FtsZ_0.17_ (WT):ΔCTL) demonstrate that increasing inter-CTL distance abolishes low repulsion and increases curvature, supporting the model that CTL-mediated electrostatics act as a tunable rheostat for Z-ring assembly and architecture.

Further comparative analysis across prokaryotic species suggests a relationship between cell dimensions and CTL charge segregation. Prokaryotes with wider cells can have low-κ CTLs, which promote inter-CTL attraction and low filament curvature, whereas high-κ CTLs, which enforce stronger curvature, are restricted to narrow bacteria. Thus, the CTL sequence encodes a simple electrostatic scaling rule that couples filament geometry to cell size.

Additionally, we find altering the CTL electrostatics *in vivo* disrupts Z-ring organization. When ΔCTL FtsZ is co-expressed with a fluorescent version of FtsZ with the wild-type CTL, rings are misaligned and deviate from the normal division plane, which impairs growth, underscoring the importance of properly tuned CTL-CTL interactions for correct division plane placement and cytokinesis.

Symmetric cell division in prokaryotes requires the Z-ring to orient precisely orthogonal to the long cell axis to ensure equal partitioning of cellular contents. While curvature-sensing mechanisms involving FtsA and FtsN have been proposed to align FtsZ filaments during later stages of division^12^, the initial cues that guide Z-ring assembly orthogonally to the long cell axis prior to FtsN recruitment remain unclear. We propose that an intrinsic match between filament curvature and the cell diameter could facilitate initial Z-ring assembly, providing an optimal starting point for divisome maturation. Because CTL-CTL interactions are weak and reversible, they would still allow for FtsZ filaments to be flexible enough allowing the Z-ring to constrict progressively as septation proceeds. In addition, the strength of the inter-CTL interactions could be dynamically modulated during cytokinesis through binding of regulatory proteins or by CTL phosphorylation^49^, which could alter net charge and charge patterning and modulate the electrostatic contacts between CTLs^50^.

To conclude, our findings suggest that FtsZ’s CTL acts as a curvature-determining element that links molecular sequence to cellular geometry, exemplifying how intrinsically disordered linkers can encode geometric information and provide bacteria with a simple, evolvable strategy for achieving precise and symmetric division.

## Acknowledgments

We thank all members of the Loose group (ISTA) for helpful discussions, particularly Dr. Benjamin Springstein and Dr. Biswaprakash Mahanta for their comments on the manuscript. We are grateful for Dr. Benjamin Springstein’s help regarding *in vivo* experiments, Roman Hajdu and the group of Andela Saric (ISTA, Austria) for helpful discussions. We would also like to thank the groups of Dr. Calin Guet (ISTA, Austria) and Dr. Andrea Vettiger (University of Lausanne, Switzerland) for gifting us *E. coli* MG1655 and *E. coli* MG1655 AV72, respectively. This research was supported by Scientific Services Units of ISTA: Image and Optics Facility, Lab Support Facility and Scientific Computing Facility.

## Author contributions

Project management and funding acquisition: M.L; Conceptualization and design: M.L and M.K; Experimental investigation: M.K; Data and image analysis: M.L, M.K and D.M; Visualization: M.L and M.K; Manuscript writing: M.L and M.K.

## Material and Methods

### Identification and analysis of Intrinsically Disordered Regions (IDRs)

A dataset of 4,603 FtsZ sequences was compiled from bacterial and archaeal species with cell width measurements annotated in the BacDive database ^27^. Intrinsically disordered regions (IDRs) within the retrieved FtsZ protein sequences were identified using three complementary approaches. The ALBATROSS^37^ Colab notebook (https://colab.research.google.com/github/holehouse-lab/ALBATROSS-colab/blob/main/idrome_constructor/idrome_constructor.ipynb) was used for sequence-based disorder prediction. This workflow employs metapredict v2.1 (https://metapredict.net) to predict residue-level disorder probabilities^30^. Predictions were independently validated using the PrDOS web server (https://prdos.hgc.jp/about.html), which integrates local amino acid sequence information and homologous structural templates to predict intrinsically disordered regions.^31^ In parallel, structural models for all FtsZ sequences were generated using AlphaFold2 via the ColabFold implementation^51^. Predictions were performed on the ISTA high-performance computing cluster. Disordered residues were defined as those with predicted local distance difference test (pLDDT) scores < 60. A gap-tolerant segmentation algorithm allowed short structured interruptions (≤4 residues) within otherwise disordered regions. In all cases, the C-terminal IDR was defined as the longest continuous disordered segment starting at or beyond residue 310. All IDRs identified by the three approaches were analyzed in the SPARROW toolkit (https://github.com/idptools/sparrow, commit version as of March 2025) to calculate sequence-derived biophysical properties, including the fraction of charged residues (FCR), charge patterning (κ), hydropathy, and net charge per residue (NCPR). Finally, from the resulting JSON score files (rank_001 only), C-terminal IDRs were extracted by scanning residues ≥310 for the longest gap-tolerant disordered segment (disorder defined as pLDDT < 60, allowing ≤4-residue structured interruptions), and exporting protein_id, idr_start, idr_length, and idr_end to a summary CSV. Structured regions within the CTL were defined as pLDDT > 80 and beyond position 310.

### Bacterial strains and growth conditions

*E. coli* DH5*α* (New England Biolabs (NEB), USA) strain was used for plasmid propagation during molecular cloning, while BL21 (DE3) (NEB, USA) strain was used for heterologous recombinant protein expression. *E. coli* MG1655 (courtesy of the Guet group, ISTA, Austria) was used for growth curve analysis, while MG1655 AV72 strain (courtesy of the Vettiger group, University of Lausanne, Switzerland), which carries chromosomally encoded FtsZ-G55-mVenus-Q56 as a sole FtsZ copy, was used for *in vivo* imaging. All strains were routinely grown in LB medium supplemented with an adequate antibiotic (Ampicillin (Amp) at 100 µg/mL and Kanamycin (Kan) at 50 µg/mL) at 37 ⁰C with shaking (250 rpm).

### Molecular cloning

To construct expression plasmids for the purification of FtsZ ΔCTL and FtsZ 25 aa, pML45-Amp (6xHis-SUMO-GCG-FtsZ WT)^2^,was PCR-amplified lacking the full CTL (316-365) region or N-terminal CTL (316-340) region, respectively. The amplicons were subjected to phosphorylation (NEB, USA) and ligation (NEB, USA), yielding pMK30-Amp (6xHis-SUMO-GCG-FtsZ ΔCTL) and pMK31-Amp (6xHis-SUMO-GCG-FtsZ 25 aa). All other CTL version genes (gBlocks purchased from IDT, USA) were cloned into modified pML45 (lacking CTL region) using the NEB assembly master mix (NEB, USA). For *in vivo* experiments, all full-length FtsZ mutant genes were PCR-amplified from their corresponding expression plasmids and cloned into linearized low expression plasmid pBAD33-Kan (courtesy of Dr. Benjamin Springstein, ISTA) also using the NEB assembly master mix (NEB, USA). All constructed plasmids (Supplementary Table 1) were propagated in DH5*α*, isolated using Monarch Plasmid MiniPrep Kit (NEB, USA) and confirmed by Sanger sequencing (Microsynth Austria GmbH, Austria). All primers used in this study are listed in Supplementary Table 2).

### Protein expression, purification and fluorescent labeling

Chemically competent *E. coli* BL21 (DE3) cells were transformed individually with expression plasmids carrying genes for all FtsZ mutants and FtsA. Overnight cultures supplemented with 100 µg/mL of Ampicillin, were diluted 1:100 times into 6 L of TB (or LB for FtsA) medium with Ampicillin (100 µg/mL). The cells were grown until they reached OD_600_ = 0.6-0.8. Then, expression was induced by addition of 1 mM isopropyl β-d-1-thiogalactopyranoside (IPTG) and cells were further incubated at 37 ⁰C for 3 h. After expression, cells were harvested by centrifugation at 6000xg for 45 min, the pellets were resuspended in the remaining supernatant, again centrifuged at 4000xg for 30 min and flash frozen in liquid N_2_. The pellets were stored at -70 ⁰C. All FtsZ mutants were purified the same way. The pellets were resuspended in Lysis Buffer (50 mM Tris-HCl pH = 8, 500 mM KCl, 20 mM imidazole, 2 mM β-mercaptoethanol and 10 % glycerol) supplemented with EDTA-free protease-inhibitor tablets (1 per 50 mL of the buffer) and 1 mg/mL DNAse I. The lysates were sonicated for no longer than 10 min (amplitude: 40, time on: 1 s, time off: 4s) using Q700 sonicator with a probe of 12.7 mm diameter immersed in the resuspension. Cell debris was further cleared by centrifugation at 31000xg for 45 min. Subsequently, cleared lysate was mixed and incubated with HisPurTM Ni-NTA resin (Thermo Fisher Scientific; 1 mL of resin per 1 L of expression culture) for 1 h at 4 ⁰C. Next, the resin was washed with 30 column volumes (CV) of Lysis Buffer and His-tagged proteins were eluted with isocratic elution using 250 mM Elution Buffers (50 mM Tris-HCl pH = 8, 500 mM KCl, 250 mM imidazole, 2 mM β-mercaptoethanol and 10 % glycerol). The fractions were pooled and dialyzed overnight at 4 ⁰C against Dialysis Buffer (50 mM Tris-HCl pH = 8, 300 mM KCl, 2 mM β-mercaptoethanol and 10 % glycerol) in the presence of His-Ulp1 protease (1:100 molar ratio) for His-SUMO cleavage. To remove cleaved His-tag and His-Ulp1, reverse affinity chromatography was applied by mixing the dialyzed protein with Protino Ni-IDA resin (LACTAN Macherey Nagel) for 30 min, after which the flowthrough was collected and analysed by SDS-PAGE. Proteins were further dialyzed into Storage buffer (50 mM Tris-HCl pH = 7.4, 50 mM KCl, 1 mM EDTA and 10 % glycerol) overnight at 4 ⁰C, after which the protein concentration was determined by Bradford Assay. At this point, all proteins were aliquoted, flash frozen in liquid N_2_ and stored at -70 ⁰C. Since all proteins were purified with Gly-Cys-Gly sequence on N-termini, were further labeled using CP400, Alexa Fluor 488 (AF488) or Atto633. The engineered Cysteine on N-terminus was used to covalently bind the fluorescent dyes. The dyes were added in 3-fold molar excess and incubated for 2 h on RT. To remove the remaining unbound dye, the samples were loaded onto PD10 desalting columns pre-equilibrated in Storage Buffer. The fractions were pooled, analyzed by SDS-PAGE and the concentration was measured by Bradford assay. Labelled proteins were stored at -70 ⁰C. FtsA was purified as previously described^52,53^. In brief, the pellet was resuspended in FtsA buffer (50 mM Tris-HCl pH = 8.0, 500 mM KCl, 10 mM MgCl_2_ and 0.5 mM DTT) with protease-inhibitor tablets (1/50 ml) and DNAse I (1 mg/ml). This was followed by sonication using a Q700 Sonicator as described above. Cell debris was centrifuged for 45 minutes at 23500xg and mixed with pre-cleaned Strep-Tactin beads for 1 h at 4 ⁰C. Using the FtsA buffer, washing was done until no proteins were washed, which was indicated by Bradford assay. The isocratic elution was triggered with the FtsA buffer supplemented with 5 mM desthiobiotin. Subsequently, eluted protein was incubated overnight at 4 ⁰C in the presence of Ulp1 (1:100 molar ratio). The cleaved protein was directly loaded onto a HiLoad 26/600 Superdex 200 SEC column and eluted with the FtsA storage buffer (50 mM Tris-HCl pH = 8.0, 500mM KCl, 10 mM MgCl_2_, 0.5 mM DTT and 10% glycerol). Collected fractions were analyzed by SDS-PAGE and pulled fractions were flash frozen and stored at -70 ⁰C.

### GTPase assay

GTPase activity of FtsZ mutants was observed by measuring released phosphate using the Malachite Green Assay Kit (Sigma-Aldrich, USA). The proteins were diluted in Reaction Buffer (50 mM Tris-HCl pH = 7.2, 150 mM KCl, 5 mM MgCl_2_), all of them at the 5 µM final concentration, incubated 30 min at RT in the presence or absence of 1 mM GTP and subsequently diluted 5 times in previously made Malachite Green Mixture (1 mL reagent A + 10 µL reagent B, according to manufacturer’s instructions). After the incubation of 30 min on RT, the absorbance was measured at 620 nm. The same was done for a phosphate standard of known concentrations (provided by the manufacturer), so that the obtained absorbance values for FtsZs could be translated to µM units of PO_4_^3-^ released, based on linear regression of the standard curve. For determining the critical concentrations (Cc), the same protocol was used except that the final protein concentrations were varied (0.2, 0.3, 0.4, 0.5, 0.6, 0.7, 0.8 and 0.9 µM). The Cc was defined as the concentration at which the PO_4_^3-^ release was significantly increased.

### Supported lipid bilayers (SLB) preparation for *in vitro* imaging

Chloroform solutions of synthetic DOPC (1,2-dioleoyl-sn-glycero-3-phosphocholine) (25 mg/ml) and DOPG (1,2-dioleoyl-sn-glycero-3-phospho-(1‘-rac-glycerol)) (25 mg/ml), were mixed in 67:33 mol% ration and dried under the N_2_ stream. Dried lipid films were rehydrated in a Swelling Buffer (50 mM Tris-HCl pH = 7.2, 300 mM KCl) to a final concentration of 5 mM, incubated at 37 ⁰C for 1 h and thoroughly vortexed. This was followed by five freeze-thaw cycles and sonication (amplitude: 1, time on: 1 s, time off: 4 s, total process time: 5 min) in order to produce Small Unilamellar Vesicles (SUV). SUV suspension was further diluted in a Swelling Buffer to 0.5 mM, mixed with 1 mM CaCl_2_ and put into a reaction chamber that is attached to a piranha-treated glass coverslip (30 % H_2_O_2_ and 98 % H_2_SO_4_ mixed in 1:3 ratio; 2 h) and incubated at 37 ⁰C for 1 h to allow for SLB formation. The remaining unfused SUVs were removed by thorough wash with 2x excess of Reaction Buffer.

### Growth curve analysis

Low expression plasmids containing FtsZ charge segregation mutants were individually transformed into chemically competent *E. coli* MG1655 cells. Single colonies were then transferred into overnight cultures, which were diluted into 100 well plates 100 times. The expression of mutants was induced with addition of 0.2 % arabinose. Growth was followed by measuring OD at 600 nm every hour while constant shaking at 37 ⁰C using the Bioscreen growth analysis system (Oy Growth curves Ab Ltd, Finland).

### Agar pad preparation for *in vivo* imaging

*E. coli* MG1655 AV72 was transformed individually with pMK69, pMK71 or pBAD33.1, single colonies were transferred into LB with 50 µg/mL kanamycin and grown overnight at 37 ⁰C with shaking. The culture was refreshed 100 times into LB (50 µg/mL kanamycin) supplemented with or without 0.2 % arabinose and grown at 37 ⁰C with shaking until the OD_600_ of 0.5 was reached. The cells were diluted 1:5 and 3 µL were plated on M9 agar pad (M9 salts + 0.2 % glucose, 2 mM MgSO_4_, 0.1 mM CaCl_2_ and 1 % low-melting agarose; with or without 0.2 % arabinose) on the glass slide, left to dry, a coverslip was put on top of the pad and sealed with VALAP (1:1:1 mixture of vaseline, lanolin, and paraffin).

### Total internal reflection fluorescence (TIRF) microscopy

The Visitron iLAS2 TIRF system had 100x Olympus TIRF NA 1.46 oil objective with Laser Quad Band Filter (405/488/561/640 nm) to filter emitted fluorescence. The signals were recorded on Photometrics Evolve 512 EMCCD (512 x 512 pixels, 16 x 16 μm^2^) operating at a frequency of 5 Hz. Dual color and *in vivo* imaging were done on a Nikon Eclipse Ti2 microscope with iLas2 1250 (GATACA) 360° Ring TIRF module, using a CFI 1242 Plan Apo Lambda D1 100x oil immersion objective (NA 1.45, WD 0.13 mm), Omicron LightHUB Ultra as a laser source, and Nikon 525/50 ET bandpass coupled to Cairn TwinCam camera splitter. Time-lapse experiments were performed with the following parameters: EM gain of 200; exposure time of 50 ms; frame interval of 2s (1s for *in vivo* imaging); total time of 15 min (10 min for *in vivo* imaging) and laser power of 4-20%. Fluorescent recovery after photobleaching (FRAP) experiments were done with the following parameters: EM gain of 200; 10 pre-FRAP number of frames; 110 post-FRAP number of frames; FRAP laser power of 40-60%; time-lapse laser power of 4-20%; frame interval of 1s; dwell size of 1 µs/pixel and overlapping lines of 75%. The Visitron iLAS2 TIRF microscope’s resolution is 0.1 µm per pixel, whilst Nikon Eclipse Ti2 microscope’s resolution is 0.06 µm per pixel.

### Image and Data analysis

Treadmilling and FRAP analysis from TIRF movies were done by in-house built scripts based on ImageJ plug-ins and Python.^40^ Represented micrographs were created by averaging 50 frames with adjusted contrast. PCC analysis of dual color images was done using the Coloc2 ImageJ plug-in. All experiments are performed at least three times (n=3) and the numeric data are written in the form of mean ± standard deviation (S.D.). For statistical analysis, we used two-tailed Student’s *t*-test and Pearson correlation test.

## Supplementary Figures and Figure Legends

**Fig. S1.**
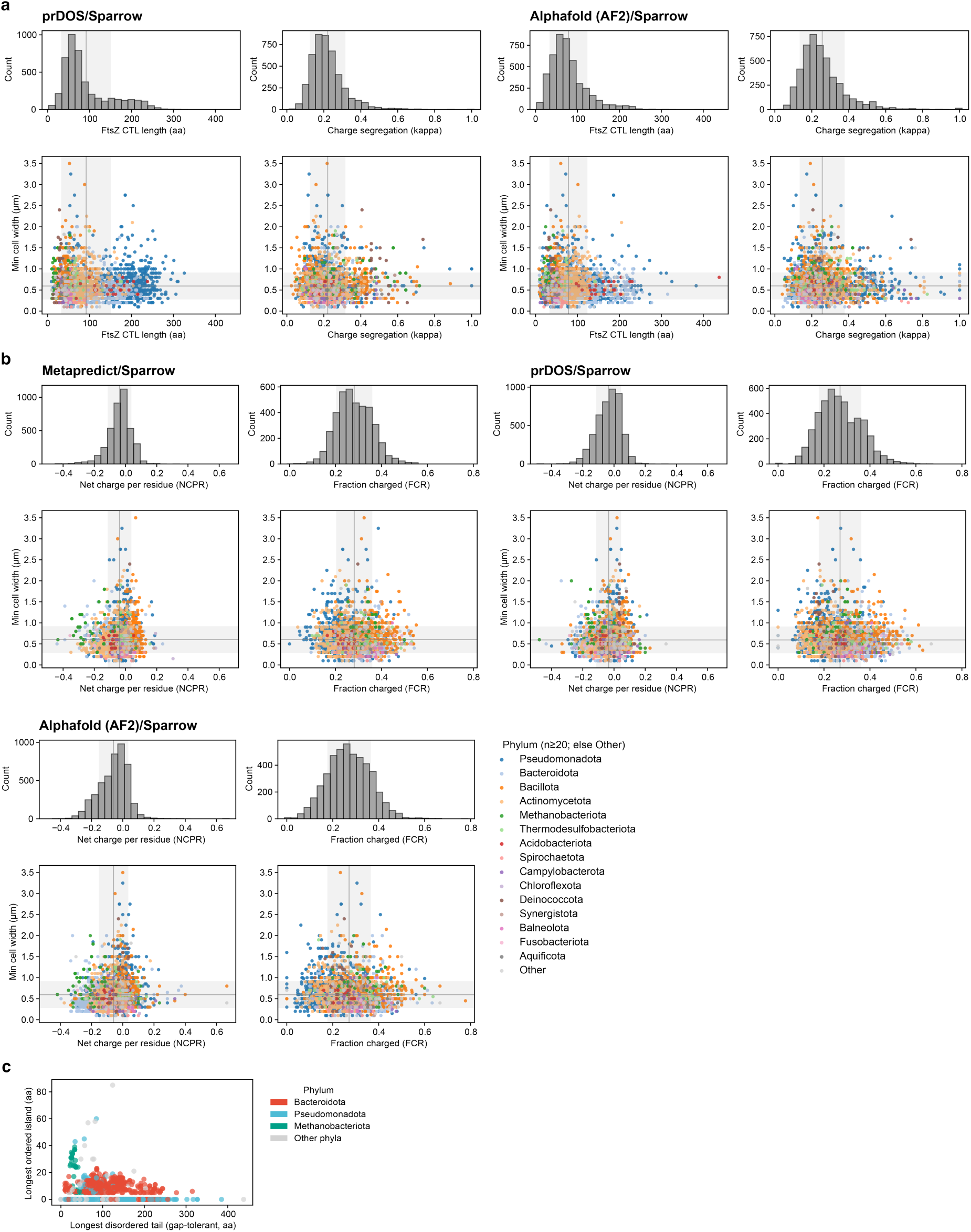
Correlations between FtsZ C-terminal linker (CTL) biophysical properties across and cell widths. **(a)** Relationship between cell widths and the length of CTL (left) or CTL’s charge segregation (right) analyzed using AlphaFold2 (left) or PrDOS-identified (right) CTLs. Different colors represent different phyla. Histograms represent the distribution of CTL lengths or charge segregation. Gray line and gray-shaded area represent the mean and standard deviation of the corresponding parameters. **(b)** Relationship between cell widths and net charge per residue (NCPR) or fraction of charged residues (FCR) using all three predictors for the identification of CTLs. Histograms represent the distribution of the corresponding parameters. **(c)** Longest ordered island vs. longest disordered tail in our dataset.

**Fig. S2.**
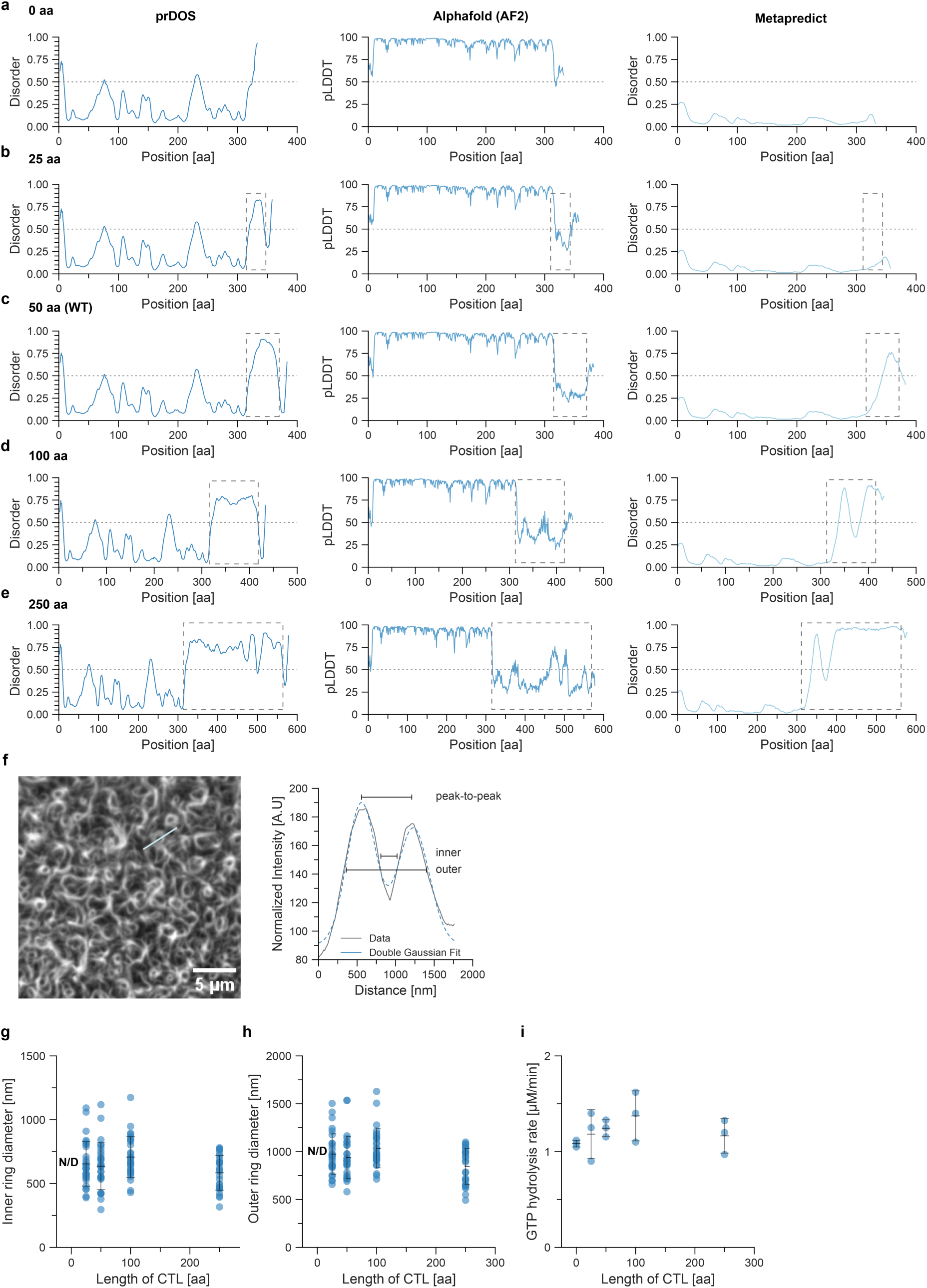
Effect of the length of CTL on large-scale self-organization of FtsZ networks. **(a-e)** Predicted intrinsic disorder profiles across CTL variants of increasing length (0 aa, 25 aa, 50 aa (WT), 100 aa, and 250 aa), as calculated using prDOS, AlphaFold-pLDDT, and Albatross. Each plot shows disorder probability as a function of residue position, demonstrating progressive increases in predicted disorder with CTL extension. **(f)** Representative normalized fluorescence intensity profile of a Z-ring, with a double Gaussian function fit used to quantify inner ring diameter, outer ring diameter, and peak-to-peak diameter (illustrated as measurements 1–3). Analysis of inner **(g)**, outer **(h)** ring diameter (nm) and GTP hydrolysis rate (protein concentration at 5 µM) **(i)** in respect to the length of CTL (Fig. 2c).

**Fig. S3.**
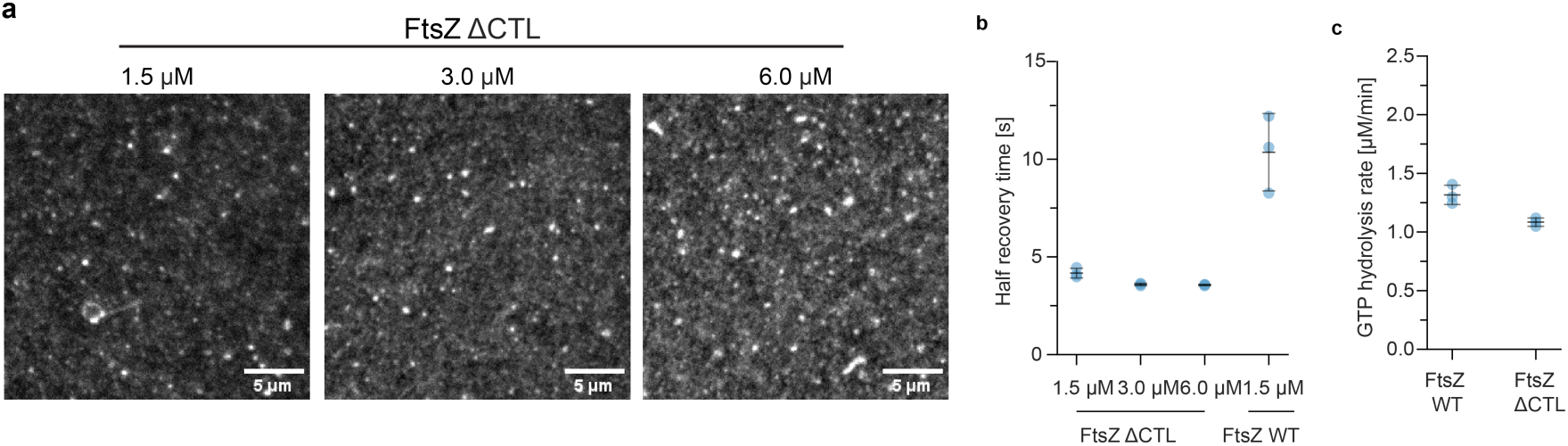
Effect of CTL deletion on large-scale self-organization of FtsZ networks. **(a)** TIRF micrographs of fluorescently labeled FtsZ ΔCTL at increasing concentrations (1.5, 3 and 6 µM) on SLBs recruited by FtsA (0.2 µM). Scale bar: 5 µm. **(b)** Analysis of half recovery time in (a) and comparison to half recovery time of FtsZ WT. **(c)** Comparison of GTP hydrolysis rates between FtsZ WT and FtsZ ΔCTL (protein concentration at 5 µM).

**Fig. S4.**
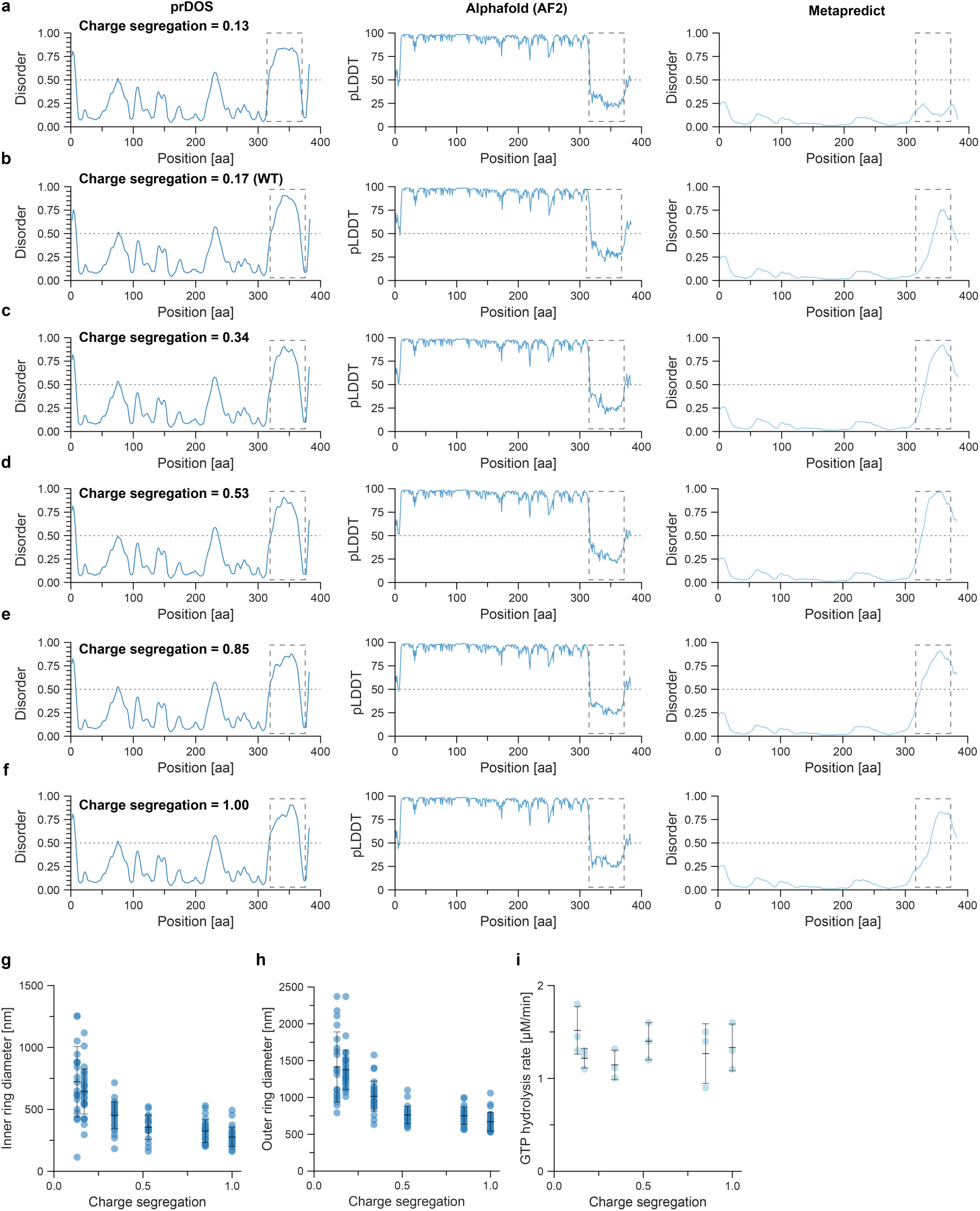
Effect of the charge segregation in CTL on large-scale self-organization of FtsZ networks. **(a-f)** Predicted intrinsic disorder profiles across CTL variants of increasing charge segregation (κ values of 0.13, 0.17 (WT), 0.34, 0.53, 0.85 and 1.00), as calculated using prDOS, AlphaFold-pLDDT, and Sparrow. Analysis of inner **(g)**, outer **(h)** ring diameter (nm) and GTP hydrolysis rate (protein concentration at 5 µM) **(i)** in respect to the charge segregation in CTL (Fig. 3b).

**Fig. S5.**
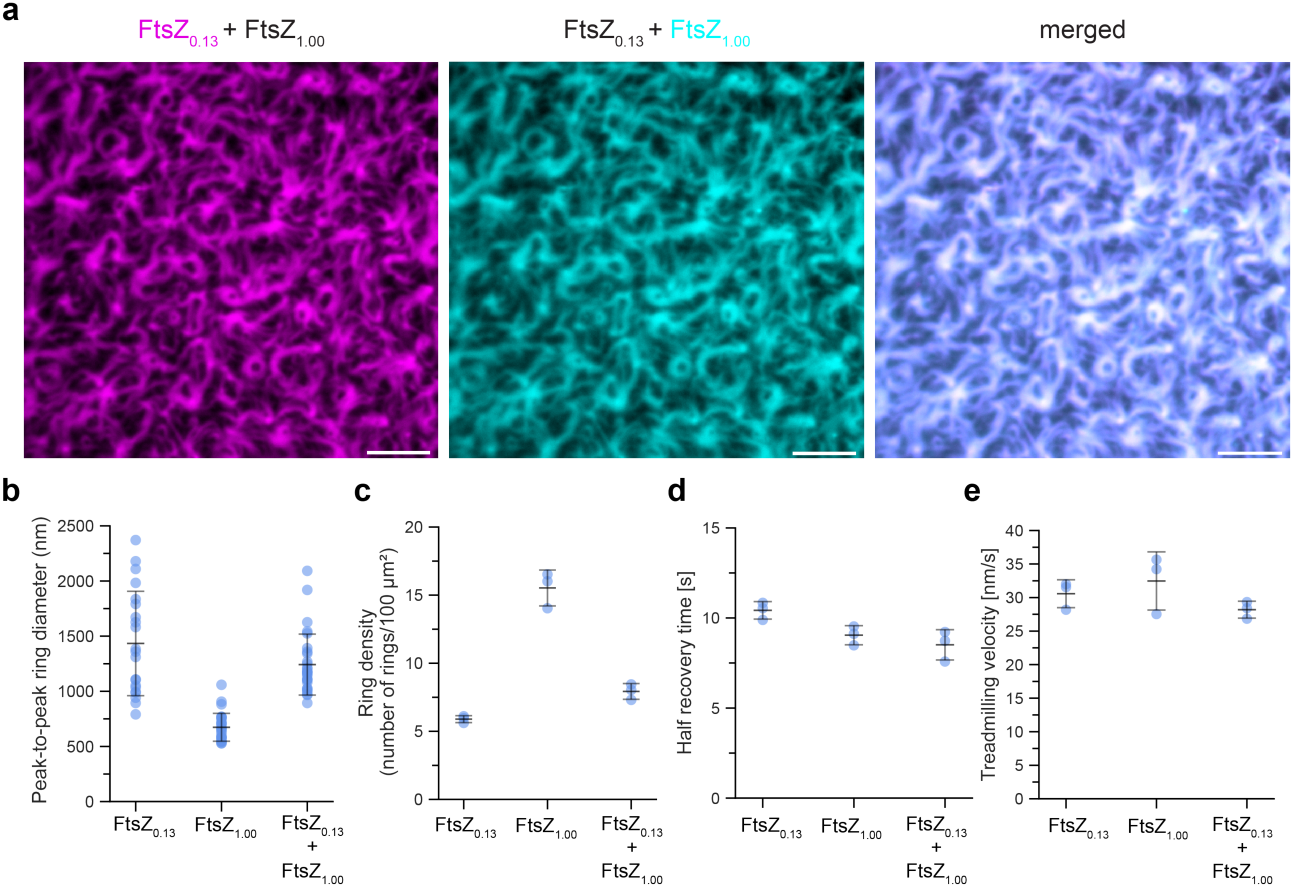
Co-assembly of mixed-charge-segregation FtsZ variants alters Z-ring architecture. **(a)** TIRF micrographs of fluorescently labeled both FtsZ_0.13_ (Atto633) and FtsZ_1.00_ (Alexa488) (1.5 µM) on SLBs recruited by FtsA (0.2 µM). Scale bar: 5 µm. **(b)** Analysis of peak-to-peak ring diameter (nm), **(c)** ring density (number of rings/100 µm^2^), **(d)** half recovery time (s) and **(e)** treadmilling velocity (nm/s) from (a).

**Fig. S6.**
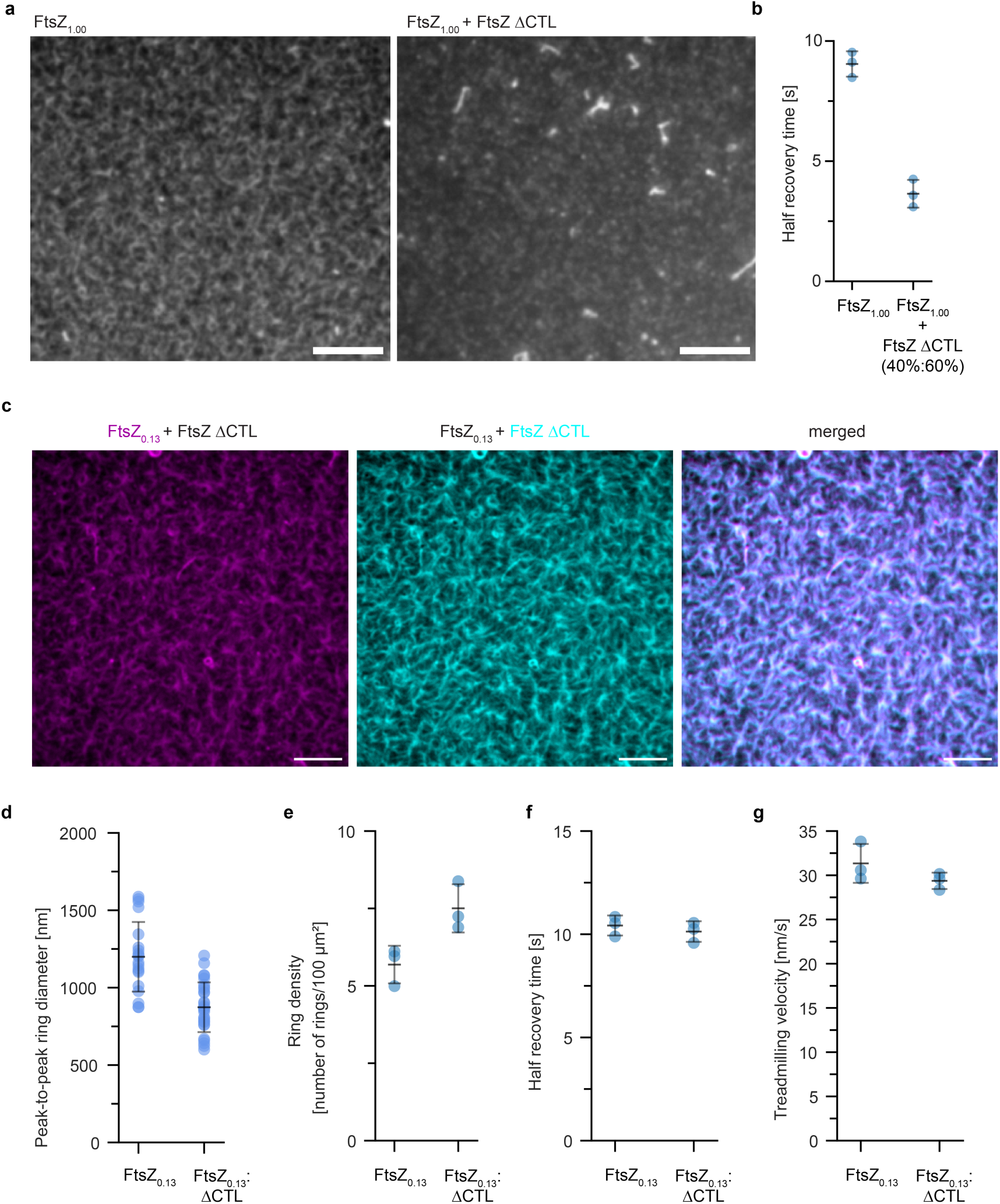
Co-assembly of full-length FtsZ variant and FtsZ ΔCTL result in formation of mixed filaments. **(a)** TIRF micrographs of Alexa488-FtsZ_1.00_ alone or mixed with FtsZ ΔCTL. Scale bar: 5 µm. **(b)** Half recovery time (s) from (a). **(c)** TIRF micrographs of fluorescently labeled both FtsZ_0.13_ (Atto633) and FtsZ ΔCTL (Alexa488) (2:3, 1.5 µM) on SLBs recruited by FtsA (0.2 µM). Scale bar: 5 µm. **(d)** Analysis of peak-to-peak ring diameter (nm), **(e)** ring density (number of rings/100 µm^2^), **(f)** half recovery time (s) and **(g)** treadmilling velocity (nm/s) from (c).

**Fig. S7.**
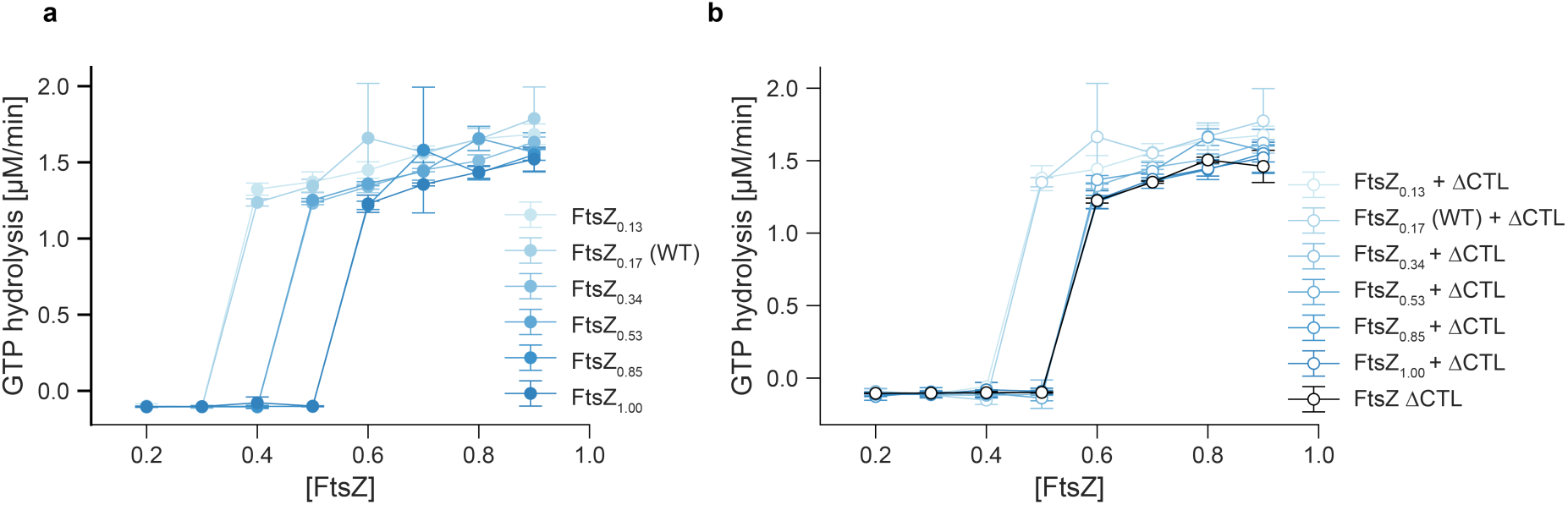
Critical GTP hydrolysis concentration (Cc) determination of the charge segregation FtsZ variants. **(a)** Critical GTPase concentration (Cc) determination of all FtsZ charge segregation variants. **(b)** Critical GTPase concentration (Cc) determination of all FtsZ charge segregation variants mixed with FtsZ ΔCTL in 2:3 ratio.

**Fig. S8.**
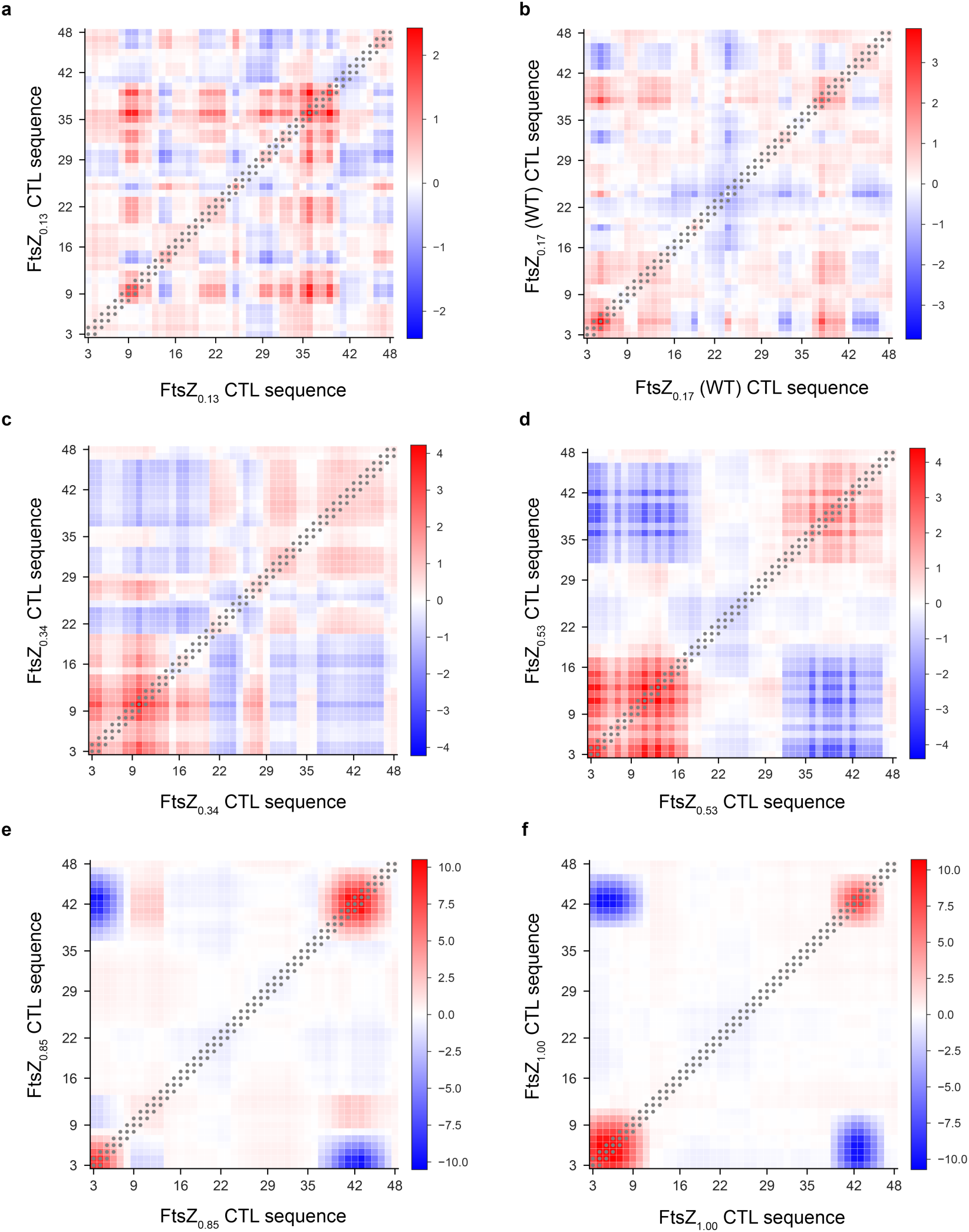
Calculation of diagonal epsilon (ε) values. **(a-f)** Intermaps of different CTLs used in this study. Gray dots indicate the values that are used to calculate mean ε with respect to configuration of CTLs in FtsZ filaments.

**Fig. S9.**
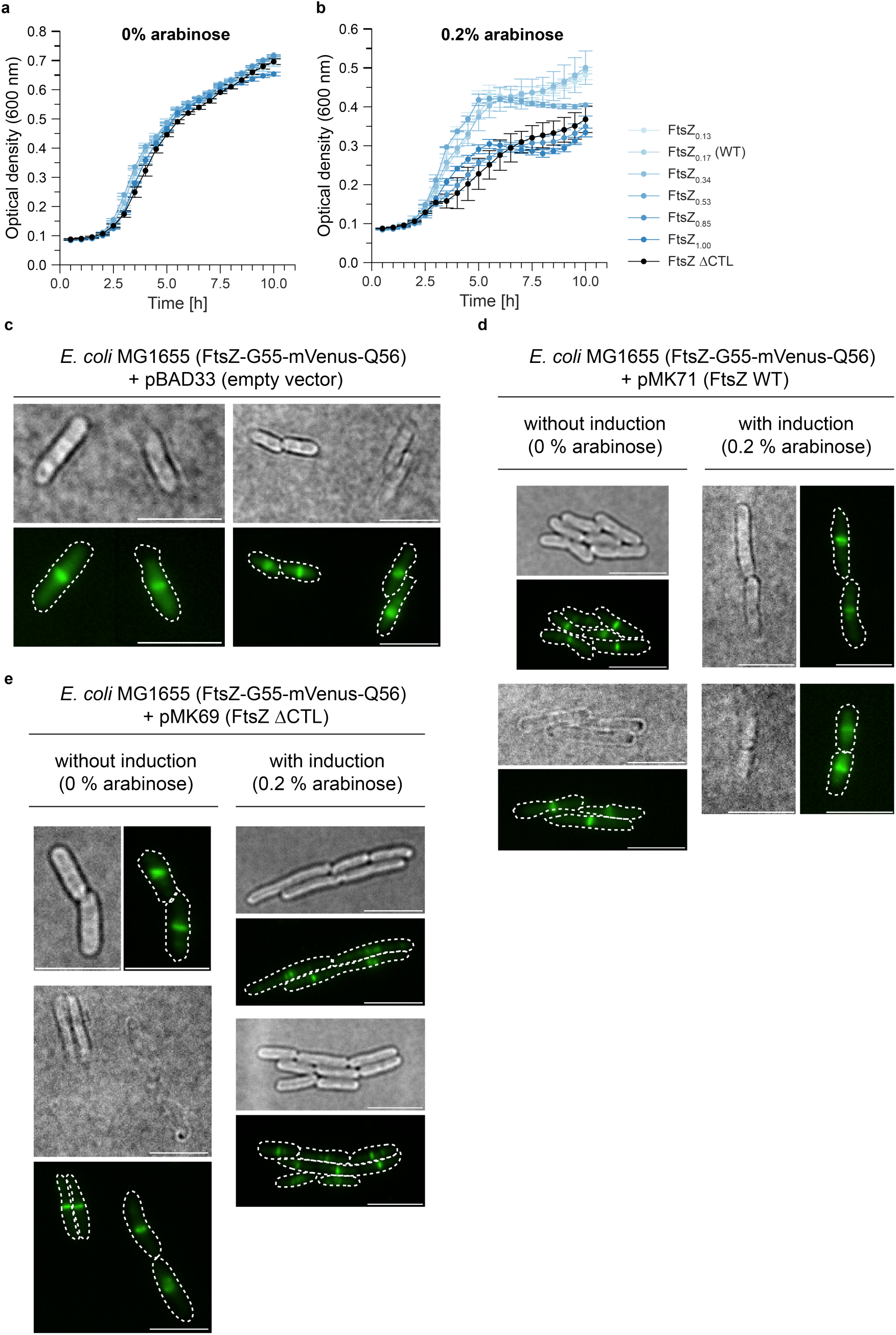
Effect of inter-CTL interactions on cell division rate *in vivo*. **(a)** Growth curve analysis (Optical Density (OD) measurement) of *E. coli* MG1655 individually transformed with plasmids carrying FtsZ charge segregation variants under non-inducible conditions (0% arabinose). **(b)** Growth curve analysis (OD measurement) of *E. coli* MG1655 individually transformed with plasmids carrying FtsZ charge segregation variants under inducible conditions (0.2% arabinose). **(c)** Direct Illumination Acquisition (DIA) and TIRF micrograph of *E. coli* MG1655 (FtsZ-G55-mVenus-Q56) transformed with pBAD33 (empty vector). Scale bar: 5 µm. **(d)** DIA and TIRF micrographs of *E. coli* MG1655 (FtsZ-G55-mVenus-Q56) transformed with pMK71 (FtsZ WT) without (left) or with (right) induced expression (0.2% arabinose). **(e)** DIA and TIRF micrographs of *E. coli* MG1655 (FtsZ-G55-mVenus-Q56) transformed with pMK69 (FtsZ ΔCTL) without (left) or with (right) induced expression (0.2% arabinose). Scale bar: 5 µm.

